# Interareal synaptic inputs underlying whisking-related activity in the primary somatosensory barrel cortex

**DOI:** 10.1101/2023.08.10.552729

**Authors:** Masahiro Kawatani, Kayo Horio, Mahito Ohkuma, Wan-Ru Li, Takayuki Yamashita

## Abstract

Body movements, especially orofacial movements, are known to influence brain-wide neuronal activity. In the sensory cortex, thalamocortical bottom-up inputs and motor-sensory top-down inputs are thought to affect the dynamics of membrane potentials (V_m_) of neurons and change their processing of sensory information during movements. However, direct perturbation of the axons projecting to the sensory cortex from other remote areas during movements has remained unassessed, and therefore the interareal circuits generating motor-related signals in sensory cortices are still unclear. Using a G_i_-coupled opsin, eOPN3, we here inhibited interareal signals incoming to the whisker primary somatosensory cortex (wS1) of awake behaving mice and tested their effects on whisking-related changes in neuronal activities in wS1. Spontaneous whisking in air induced the changes in spike rates of a fraction of wS1 neurons which were accompanied by depolarization and substantial reduction of slow-wave oscillatory fluctuations of V_m_. Despite an extensive innervation, inhibition of inputs from the whisker primary motor cortex (wM1) to wS1 did not alter the spike rates and V_m_ dynamics of wS1 neurons during whisking. In contrast, inhibition of axons from the whisker-related thalamus (wTLM) and the whisker secondary somatosensory cortex (wS2) to wS1 largely attenuated the whisking-related supra- and sub-threshold V_m_ dynamics of wS1 neurons. Our findings thus suggest that sensorimotor integration in wS1 during spontaneous whisking is mediated by direct synaptic inputs from wTLM and wS2 rather than from wM1.

**Significance statement:** The traditional viewpoint underscores the importance of motor-sensory projections in shaping movement-induced neuronal activity within sensory cortices. However, this study challenges such established views. We reveal that the synaptic inputs from the whisker primary motor cortex do not alter the dynamics of neuronal activity in the whisker primary somatosensory cortex (wS1) during spontaneous whisker movements. Furthermore, we make a novel observation that inhibiting inputs from the whisker secondary somatosensory cortex (wS2) substantially curtails movement-related activities in wS1. These findings provoke a reconsideration of the role of motor-sensory projections in sensorimotor integration and bring to light a new function for wS2-to-wS1 projections.

## Introduction

Classical research on anesthetized animals demonstrated that neuronal activity in primary sensory cortices is predominantly shaped by sensory inputs (Mountcastle et al., 1957; Katsuki et al., 1959; Hubel and Wiesel, 1962). However, more recent studies conducted with awake animals have established that spontaneous behaviors such as locomotion and whisking substantially modulate sensory responses of neurons in the primary visual, auditory, and somatosensory areas (Gallant et al., 1998; Crochet and Petersen, 2006; Ferezou et al., 2006; Niell and Stryker, 2010; Polack et al., 2013; Schneider et al., 2014; Zhou et al., 2014). Although these findings highlight the functional importance of motor-related modulation in the sensory cortex, the neuronal circuit mechanisms underlying these effects are still not fully understood.

The whisker primary somatosensory barrel cortex (wS1) in rodents is among the most extensively studied sensory areas in awake mammalian brains. When mice actively explore their surroundings by rhythmic whisking, wS1 layer 2/3 (L2/3) excitatory neurons slightly depolarize and significantly reduce slow-wave fluctuations of membrane potentials (V_m_) (Crochet and Petersen, 2006; Poulet and Petersen, 2008). L2/3 neurons of wS1 exhibit sparse firing properties, and whisking-related signals may not directly influence their firing rates (Crochet and Petersen, 2006; Poulet and Petersen, 2008; Yamashita et al., 2013). However, neurons in deeper layers exhibit higher rates of spontaneous firing, and spikes of a subset of layer 5 or 6 (L5/6) neurons accurately reflect various variables of whisker movements (Fee et al., 1997; de Kock and Sakmann, 2009; Cheung et al., 2020; Suryadeep et al., 2022). Thalamocortical signals (Poulet et al., 2012), cholinergic modulation (Eggermann et al., 2014; Gasselin et al., 2021), and top-down inputs from the whisker primary motor cortex (wM1) (Lee et al., 2013; Zagha et al., 2013; Naskar et al., 2021) are suggested to play a role in generating supra- and sub-threshold V_m_ dynamics of wS1 neurons during whisking.

The involvement of interareal inputs in whisking-related activity in wS1 has mainly been inferred from the inhibition of source regions such as the whisker-related thalamic region (wTLM) (Poulet et al., 2012) and wM1 (Xu et al., 2012; Lee et al., 2013). Regional inhibition experiments could involve multiple interconnected brain regions (Otchy et al., 2015). Therefore, direct perturbation of axonal terminals within wS1 would be necessary to examine the role of monosynaptic connections between wS1 and other brain regions. Some previous studies that involve optogenetic stimulation of specific input pathways into sensory cortices revealed the existence of mechanisms that can mimic movement-related state changes (Zagha et al., 2013; Schneider et al., 2014). However, there have been no studies that include specific inhibition of synaptic inputs incoming to primary sensory cortices during motor-related activities. Furthermore, most findings have focused on the superficial layers, leaving the effects of perturbation on the deeper layers largely unexplored.

In this study, we investigate the specific contributions of different inputs to the whisking-related activity of wS1 neurons using awake head-restrained mice. We utilize an inhibitory G protein-coupled receptor (GPCR) opsin, eOPN3, to inactivate axon terminal activity in three regions that provide input to wS1 (Mahn et al., 2021): wTLM, wM1, and the whisker secondary somatosensory cortex (wS2). We first confirmed that photo-inhibition through eOPN3-mediated mechanisms is effective *in vitro* and *in vivo*. We then examined the role of each projection to wS1 in spontaneous and whisking-induced spiking activities and subthreshold V_m_ in wS1 neurons. Our findings suggest that wM1→wS1 signaling does not significantly influence whisking-induced wS1 dynamics. Instead, we demonstrate that wTLM→wS1 and wS2→wS1 signaling contribute substantially to the whisking-induced increase in spike rates. Our results redefine the understanding of sensorimotor integration in wS1 during spontaneous whisking, highlighting the significance of inputs from wTLM and wS2 but not wM1 as previously assumed.

## Materials and Methods

### Animals

All procedures for animal experiments followed the guidelines of the Physiological Society of Japan and were approved by the Institutional Animal Care and Use Committee of Fujita Health University. Adult male C57BL6/J mice (2-6 months) were used for experiments. Mice were housed in a temperature-controlled room (23 ± 2 °C) with 12 hours light/dark cycle (light 0 p.m. to 0 a.m.), with free access to food and water. All experiments using live mice were conducted in the dark phase.

### Viral production

For the production of an adeno-associated virus (AAV) vector introducing eOPN3 expression, HEK293 cells were transfected with a plasmid encoding eOPN3 (pAAV-CaMKIIa(0.4)-eOPN3-mScarlet-WPRE, purchased from Addgene, #125712) together with pHelper and pAAV-RC (serotype 9), using a standard calcium phosphate method. After three days, transfected cells were collected and suspended in lysis buffer (150 mM NaCl, 20 mM Tris pH 8.0). After four freeze-thaw cycles, the cell lysate was treated with 250 U/ml benzonase nuclease (Merck) at 37 °C for 10 − 15 min with adding 1 mM MgCl_2_ and then centrifuged at 4°C at 1753×*g* for 20 min. AAV was then purified from the supernatant by iodixanol gradient ultracentrifugation. The purified AAV solution was concentrated in PBS via filtration and stored at −80°C.

### Animal preparation and surgery

Mice were anesthetized with isoflurane (2–3% for induction, 1.0–1.5% for maintenance) and head-fixed on a stereotaxic device. Body temperature was maintained at ∼37°C by a controlled heating pad. An ocular ointment was applied over the eyes to prevent drying. AAV-CaMK2a(0.4)-eOPN3-mScarlet-WPRE (the original titer: 4.87×10^12^ copies/ml, diluted to 1/2) or AAV-Ef1a-mCherry (#114470-AAV9, obtained from Addgene; titer: 1.0×10^13^ copies/ml, diluted to 1/6) were injected into wM1 (AP: +1.0 mm, ML: −1.0 mm, DV: 350 and 850 μm), wTLM (AP: −1.8 mm, ML: −1.8 mm, DV: −3.0 mm) or wS2 (identified by intrinsic imaging) of the left hemisphere. Total injection volume was 400 nl for wM1, 100−120 nl for wS2, and 200-400 nl for wTLM. After the AAV injection, mice were kept in a home cage for at least five weeks before experiments.

Prior to the recording, mice were implanted with a light-weight metal head holder and a recording chamber under isoflurane anesthesia (Yamashita et al., 2013). The chamber was made on the left hemisphere by building a thin wall with dental cement, and the exposed skull was covered by a thin layer of dental cement. We also implanted a small metal screw over the cerebellum as a reference electrode for silicon probe recordings. The locations of the C2 barrel column and wS2 were identified through intrinsic optical imaging (Ferezou et al., 2007; Yamashita et al., 2013) under light isoflurane anesthesia. Before recordings, mice were adapted to head restraint on the recording setup through two or three habituation sessions within two or three days. In the case of two habituation sessions, the first was 15 min, and the second was 60 min. In the case of three habituation sessions, the first session was 15 min, the second was 30 min, and the third was 60 min. On or one day before the experimental day, a small craniotomy was made over the C2 barrel column. The skull near the craniotomy was thinned for whole-cell recordings to place an optical fiber. When the mice were kept overnight after craniotomy, the exposed brain was protected with a silicon elastomer (Kwik-Cast, WPI). All the whiskers except the left and right C2 were trimmed before experiments.

### Silicon probe recording

Local field potentials (LFP) and extracellular spikes were recorded in head-fixed mice using a silicon probe (A1x32-Poly2-10mm50s-177, NeuroNexus) with 32 recording sites along a single shank covering 775 μm of the cortical depth. The probe was stained by dipping it into a DiO solution (1-2 % in ethanol) to label the recording site. The probe was then lowered gradually until the tip was positioned at a depth of 800–1100 μm under the pial surface. The craniotomy site was then covered with Ringer’s solution containing 1.5% agarose, which was further covered by paraffin to avoid desiccation of the solution. Neural data were filtered between 0.1 Hz and 7603.2 Hz, amplified using a digital head-stage (RHD2132, Intan Technologies), and digitized with a sampling frequency of 30 kHz. The digitized neural signal was transferred to an acquisition board (Open Ephys) and stored on an internal HDD of the host PC for offline analysis. To estimate the depth of L4 in each silicon probe recording, we presented a piezo sensor within reach of the right C2 whisker and recorded LFP upon active whisker touches to the sensor. The TTL pulses for the video acquisition timing, light stimulation, and piezo signals were also recorded through the Open Ephys acquisition board.

### *In vivo* whole-cell recording

Whole-cell patch-clamp recordings targeted to neurons in the C2 barrel column of awake head-restrained mice were performed using a patch-clamp amplifier (Multiclamp 700B, Molecular Devices) (Yamashita et al., 2013; Yamashita and Petersen, 2016; Wang et al., 2020). Recording pipettes (5–8 MΩ) were pulled from a borosilicate glass capillary and filled with a solution containing (in mM): 135 potassium gluconate, 4 KCl, 10 HEPES, 10 sodium phosphocreatine, 4 Mg-ATP, 0.3 Na-GTP (pH 7.3, adjusted with KOH). The recording pipettes were advanced with positive pressure (20–35 mbar), and a brief negative pressure was applied to establish a giga-ohm seal at encountered cells. All recordings were conducted under the current-clamp configuration. Liquid junction potential was not corrected.

### *In vitro* whole-cell recording

Mice were transcardially perfused under isoflurane anesthesia with an ice-cold dissection buffer containing (in mM): 87 NaCl, 25 NaHCO_3_, 25 D-glucose, 2.5 KCl, 1.25 NaH_2_PO_4_, 0.5 CaCl_2_, 7 MgCl_2_, and 75 sucrose, aerated with 95% O_2_ + 5% CO_2_. The mice were then decapitated, and the brain was isolated and cut into coronal slices (250–300 μm thick) on a vibratome in the ice-cold dissection buffer. The slices containing wM1 were incubated for 30 min at 35°C in the dissection buffer and maintained thereafter at room temperature (RT) in an artificial cerebrospinal fluid (aCSF) containing (in mM): 125 NaCl, 25 NaHCO_3_, 25 D-glucose, 2.5 KCl, 1.25 NaH_2_PO_4_, 1 MgCl_2_, and 2 CaCl_2_, aerated with 95% O_2_ and 5% CO_2_.

Whole-cell patch-clamp recordings were performed using an IPA amplifier (Sutter Instruments) at RT. Fluorescently labeled cells were visually identified using an upright microscope (BX51WI; Olympus) equipped with a scientific complementary metal-oxide-semiconductor (sCMOS) video camera (Zyla4.2plus, Andor). The recording pipettes were filled with the intracellular solution containing (in mM): 135 potassium gluconate, 4 KCl, 4 Mg-ATP, 10 Na_2_-phosphocreatine, 0.3 Na-GTP, and 10 HEPES (pH 7.3, 295 mOsm). Patch pipettes (5–7 MΩ) had a series resistance of 6.5–30 MΩ. Data were filtered at 5 kHz, digitized at 10 kHz, and recorded using the SutterPatch software running on Igor Pro 8.

### Filming whisker movement

The mouse orofacial part, including the right C2 whisker, was filmed from a top-view at 500 frames per sec using a high-speed camera (HAS-L1, Ditect). Each recording sweep was 60–75 s, synchronized to the electrophysiological recording through TTL pulses.

### Optogenetics

Photo-stimulation during *in vivo* electrophysiological recordings was conducted by application of green LED light (530 nm; M530F2, Thorlabs) through an optic fiber cannula with a fiber diameter of 400 μm and a numerical aperture of 0.39 (CFML14L05, Thorlabs). The light power at the fiber tip was measured as 7.3 mW. For silicon probe recordings, the tip of the fiber cannula was placed on the brain surface over the recording site. For whole-cell recordings, the tip of the fiber cannula was placed on the thinned skull close to the craniotomy. The green LED was turned on at 5 s before starting a whisker filming epoch and turned off at the end of the epoch. Each epoch was separated for more than 1 min. For photo-stimulation during *in vitro* whole-cell recordings, the green LED light was applied through the objective lens with a light power of 1 mW/mm^2^.

### Data analysis

#### Analysis of whisker movement

Movement of the right C2 whisker was quantified offline with ImageJ using an open-source macro (https://github.com/tarokiritani/WhiskerTracking) (Yamashita et al., 2013). Based on the whisker behavior, recording segments with one-, two, or three-second time windows were classified as a quiet (with no whisker movement) or whisking (with continuous rhythmic whisker movement) epoch. The onset of the whisking epoch was taken as the time the whisker angle exceeded 1 deg above baseline.

#### Analysis of spikes

Spiking activity recorded by a silicon probe was detected and sorted into clusters using Kilosort 2 (https://github.com/MouseLand/Kilosort/releases/tag/v2.0). After an automated clustering step, clusters were manually sorted and refined using Phy 2 (https://github.com/cortex-lab/phy). Well-isolated single units (642 units from wS1 and 55 units from wM1) were included in the dataset. These units were classified as fast-spiking (FS) putative interneurons or regular-spiking (RS) putative pyramidal cells based on their trough-to-peak time of average spike waveform. Single units with a trough-to-peak time < 0.4 ms were classified as FS cells, and units with a trough-to-peak time > 0.5 ms were classified as RS cells. Units showing an intermediate (0.4–0.5 ms) trough-to-peak time were excluded from the analysis. For classifying units based on the modulation by whisking, we compared their spike rates during quiet epochs and those during whisking epochs. The units that exhibited significantly higher or lower firing rates during whisking than during quiet epochs were classified as W-Up or W-Down units, respectively. The units showing no significant change in their firing rate upon whisking are classified as NM (non-modulated) units. To normalize spike rates for each unit, we obtained the mean and standard deviation (SD) of spike rates of 2-s bins of all filmed sweeps and calculated Z-scores. For the presentation of the Z-scored peri-event time histogram (Fig. 2*E,J*, 4*D*, 6*D*), the Z-scored spike rates of 10-ms bins were Gaussian-filtered and subtracted by the baseline (1.0-0.2 s before the whisking onset).

#### Defining layer 4 by current source density analysis

The averaged LFP signals at around active touch contacts were down-sampled to 3 kHz and low-pass filtered at 100 Hz. The LFP signals from odd or even channel numbers were separately used for further analysis. The second spatial derivative of the average LFP signals was used as current source density (CSD) (Freeman and Nicholson, 1975). Sinks were expressed as negative values, and sources were expressed as positive values. The channel with (1) the shortest peak latency for the average touch response in the LFP signal, (2) the largest peak amplitude of the average touch responses in the LFP signal, and (3) the fastest CSD sink onset time upon touches, across 16 channels (the channels either with odd or even numbers among 32 channels), was identified as a center of L4 (Pala and Stanley, 2022), which was estimated as at a depth of 500 um. Layer boundaries were defined using values reported previously (Lefort et al., 2009).

#### Analysis of membrane potentials

For whole-cell recording data, the mean and variance of V_m_ and average spontaneous AP rates were calculated in each whisking or quiet epoch and averaged in either Light ON or Light OFF trials. Fast Fourier transforms (FFTs) were computed as magnitudes for 2-s segments of the recordings. The amplitude of low-frequency V_m_ fluctuations was calculated by integrating the computed FFT at 1–5 Hz.

### Statistics

All values are expressed as box plots in the figures. The edges of the box indicate 25th and 75th percentiles and the central line indicates median value. The whiskers extend to the most extreme data points, excluding outliers. Statistical tests were performed using MATLAB or GraphPad Prism. The normality of data distribution was routinely tested. Analyses of two sample comparisons were performed using paired *t* tests when each sample was normally distributed, or using Wilcoxon rank sum (unpaired) or Wilcoxon signed rank (paired) test when at least one of the samples in every two-sample comparison was not normally distributed. When we performed multiple two-sample comparisons for the classification of W-Up, W-Down, and NM units, *p*-values were corrected using the Benjamini-Hochberg procedure to control the false discovery rate (Benjamini and Hochberg, 1995), unless otherwise noted.

## Results

### Impact of motor-sensory signaling on wS1 activity

Neuronal activity in wM1 can influence wS1 activity (Xu et al., 2012; Lee et al., 2013; Miguel et al., 2013; Zhang and Zagha, 2023). This impact can either originate directly from axonal terminals of wM1 neurons that project to wS1 (Petreanu et al., 2009; Petreanu et al., 2012; Lee et al., 2013; Zagha et al., 2013; Kinnischtzke et al., 2014; Naskar et al., 2021) or indirectly through signaling from other brain areas (Urbain and Deschênes, 2007a; Miguel et al., 2013). To elucidate the circuit mechanisms underlying wM1→wS1 signaling, we employed a G_i_-coupled inhibitory opsin, eOPN3 (Mahn et al., 2021), for optogenetic inhibition of ipsilateral wM1 or wM1 axonal projections in wS1 (wM1→wS1 axons). We virally introduced expression of eOPN3 in wM1 neurons and used the mice after a minimum of five weeks following the viral injection. We first evaluated the inhibitory effect of light-induced activation of eOPN3 on neuronal activity. In acute brain slice preparations, green LED illumination (530 nm, 1.0 mW/mm²) induced a robust hyperpolarization in eOPN3-expressing wM1 neurons (−8.0 ± 2.0 mV, *n* = 10 cells, Fig. 1*A,B*) with a mean peak latency of 8.0 ± 2.0 s (*n* = 10 cells, Fig. 1*C*). In awake head-restrained mice, green light exposure (530 nm, 7.3 mW at the fiber tip) applied to the surface of wM1 markedly reduced action potential (AP) rates of wM1 cells as recorded by a silicon probe (by 95.1 ± 1.3 %, *n* = 55 cells, Fig. 1*D-E*). We observed these photo-inhibitory effects consistently across all cortical layers (Fig. 1*F*), suggesting the efficient inhibition of neural activity *in vivo* via eOPN3-mediated mechanisms.

**Figure 1.**
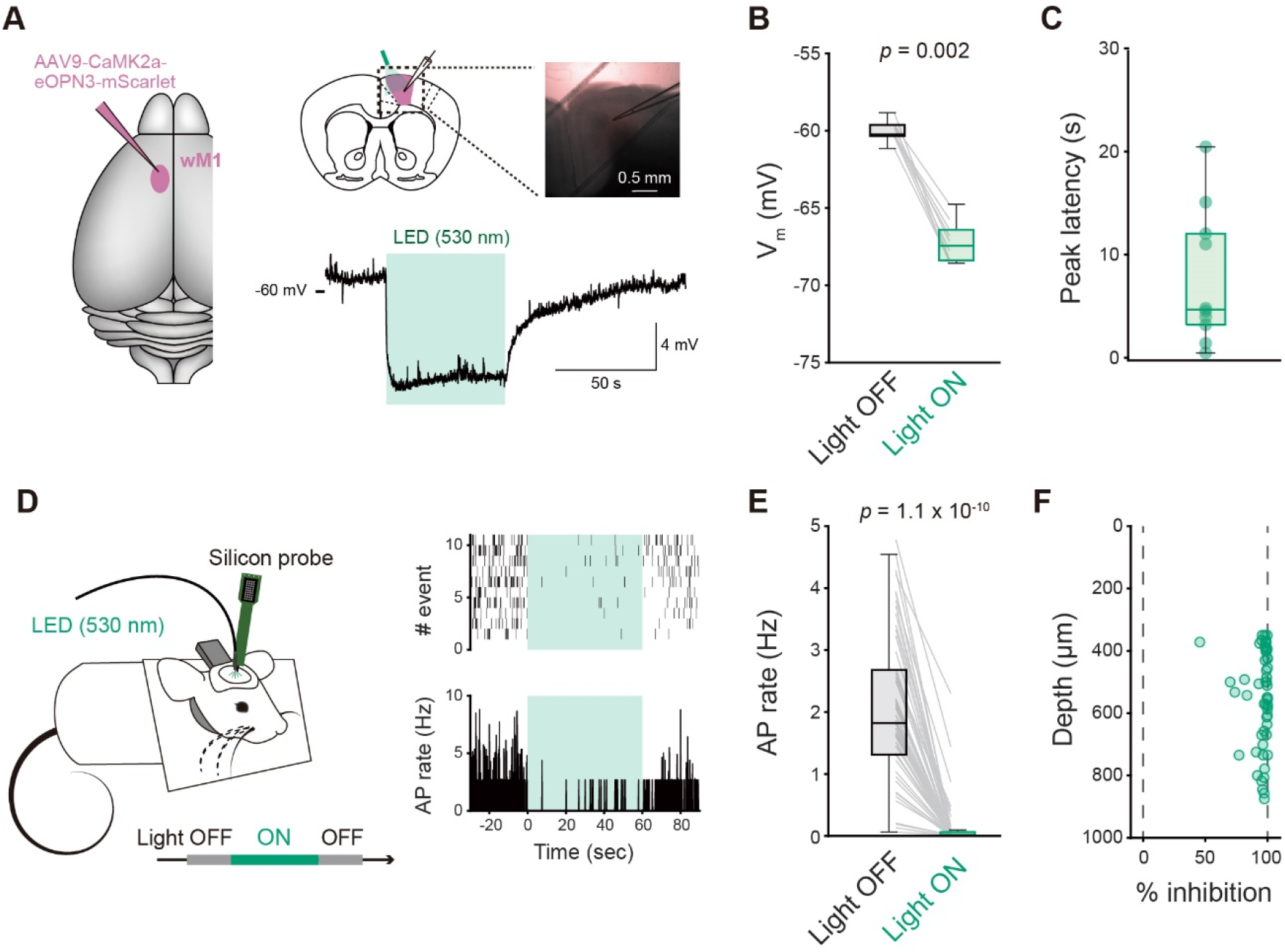
Inhibitory effect of light-induced activation of eOPN3 on neuronal activity. ***A***, Left: Schematic of viral injection. Right: Schematic of *in vitro* patch-clamp recordings (top), a bright field image of a brain slice during recording (inset) and a representative V_m_ trace upon green light illumination (bottom). Purple: eOPN3-mScarlet, green shadow: photo-stimulation. ***B***, Averaged V_m_ with (Light ON) or without (Light OFF) photo-stimulation. ***C***, Peak latencies of the responses to the photo-stimulation. ***D***, Left: Schematic of silicon probe recording from wM1 using an awake head-restrained mouse. Right: Example raster plot obtained from a representative wM1 neuron (top) and corresponding peri-stimulus time histogram (PSTH) upon green light illumination onto wM1 (green shadow). ***E***, Spontaneous AP rates of wM1 neurons with (Light ON) or without (Light OFF) green light illumination onto wM1 *in vivo*. ***F***, Percentage of the inhibition of spontaneous AP generation in wM1 plotted as a function of recording depth. *P*-values are indicated in the figure panels. Wilcoxon signed rank test (***B***, ***E***).

We next utilized these mice expressing eOPN3 in wM1 neurons for silicon probe recording from an identified left C2 barrel column in wS1 while either ipsilateral wM1 or wM1→wS1 axons in wS1 were photo-inhibited using green light (Fig. 2*A.B*). The mice were awake and head-restrained, and we filmed whisker movements at a rate of 500 Hz using a high-speed camera to compare non-whisking quiet wakefulness and the whisking state. As control experiments, we virally introduced mCherry expression in wM1 and recorded from ipsilateral wS1 while illuminating wM1 (the viral injection site) or wS1 (the recording site) with green light. During quiet wakefulness, neither photo-inhibition of wM1 → wS1 axons nor of wM1 neurons resulted in significant changes in the overall spontaneous AP rate of regular-spiking (RS) and fast-spiking (FS) neurons in wS1 compared to the corresponding control groups (Fig. 2*C,D*).

We next analyzed whisking-related neural activity in wS1 and examined the effects of photo-inhibition of wM1→wS1 axons. Among 82 RS neurons we studied, whisking induced showed an increase of AP rates in 17 neurons (20.7%, termed W-Up) without any optogenetic perturbation (Light OFF condition; Fig. 2*E,F*). In contrast, 40 (48.8%) of RS neurons exhibited a decrease (W-Down) and 25 (30.5%) showed no changes (non-modulated [NM]) in AP rates during whisking (Fig. 2*E,F*). When we photo-inhibited wM1→wS1 axons with eOPN3 (Light ON condition), 23.5 % (4 out of 17) of W-Up and 32.5% (13 out of 40) of W-Down RS units lost their responsiveness to whisking (Fig. 2*E,F*). However, the whisking-related changes in the Z-scored AP rates (W-ΔAP) in these three categories were not significantly affected by photo-inhibition of wM1→wS1 axons (W-Up: Light OFF 1.23 ± 0.13, Light ON 0.99 ± 0.18, *p* = 0.21; W-down: Light OFF -0.74 ± 0.04 Light ON -0.66 ± 0.06, *p* = 0.34; NM: Light OFF 0.09± 0.05, Light ON 0.05 ± 0.08, *p* = 0.72; Wilcoxon signed rank test for all three comparisons; Fig. 2*G*). We also analyzed FS neurons (*n* = 22 cells from 3 mice). Our recordings identified 11 (50.0%) W-Up, 5 (22.7%) W-Down, and 6 (27.3%) NM neurons among FS neurons (Fig. 2*E,F*). None of these neurons changed their category under photo-inhibition of wM1 → wS1 axons (Fig. 2*E*,*F*). The axonal inhibition did not significantly impact the W-ΔAP of FS neurons (W-Up: Light OFF 1.60 ± 0.21, Light ON 1.45 ± 0.18, *p* = 0.12; W-down: Light OFF -1.11 ± 0.15 Light ON -1.05 ± 0.22, *p* = 1.0; NM: Light OFF 0.02± 0.15, Light ON -0.22 ± 0.16, *p* = 0.10; Wilcoxon signed rank test for all three comparisons; Fig. 2*G*). These results suggest that wM1→wS1 inputs do not substantially affect the overall dynamics of wS1 activity during whisking, although they could influence the whisking responsiveness of a subset of RS neurons.

**Figure 2.**
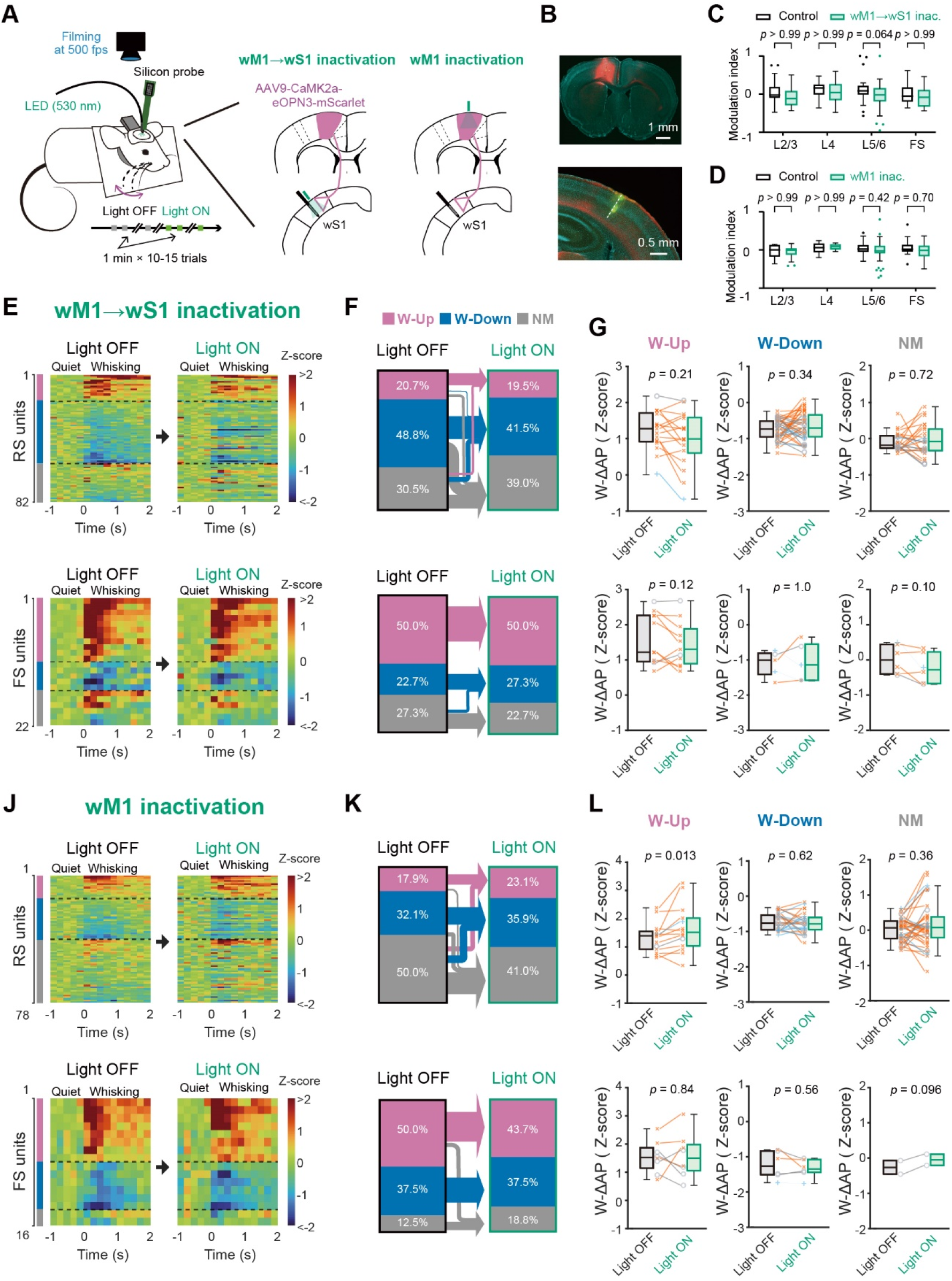
Effects of inactivation of wM1 or wM1→ wS1 axons on neuronal activity in wS1. ***A***, Schematic of the experiment. ***B***, Epifluorescence image of a coronal brain section containing the wM1 injection site (top) or the wS1 recording site (bottom). The dashed line indicates the inserted electrode’s trace. Green: DiO, red: eOPN3-mScarlet, blue: DAPI. ***C***, Modulation indices of spontaneous AP rates during quiet wakefulness upon green light illumination onto wS1 in mCherry-expressing control mice (88 cells from 2 mice) or eOPN3-expressing mice (wM1→wS1 inact., 104 cells from 3 mice). The modulation index is calculated as (AP rate in Light ON – AP rate in Light OFF)/ (AP rate in Light ON + AP rate in Light OFF). ***D***, Same as ***C***, but for photo-stimulation of wM1 (Control: 71 cells from 2 mice; eOPN3-expressing mice [wM1 inact.]: 94 cells from 2 mice). ***E***, The z-scored PSTHs of individual RS (top) and FS (bottom) units (200 ms bin size) aligned to the whisking onset with (Light ON) and without (Light OFF) inactivation of the wM1→wS1 axons. ***F***, Proportions of whisking-modulated RS (top) or FS (bottom) units in wS1 with (Light ON) and without (Light OFF) the wM1→wS1 inactivation. Purple: Positively modulated units (W-Up), blue: Negatively modulated units (W-Down), gray: Non-modulated units (NM). Arrows show transitions of whisking-modulated units from Light OFF to Light ON trials. ***G***, Whisking-related changes in the Z-scored AP rates (W-ΔAP) of RS (top) and FS (bottom) units with (Light ON) and without (Light OFF) the wM1→wS1 inactivation. ***J***–***L***, Same as ***E***–***G***, but for wM1 inactivation. Blue lines and crosses: L2/3 units, gray lines and circles: L4 units, orange lines and saltires: L5/6 units. *P*-values are indicated in the figure panels. Bonferroni’s multiple comparison test (***C***, ***D***), Wilcoxon signed rank test (G: RS W-Up, RS W-Down, RS NM, FS W-Down and FS W-Up, L: RS W-Up, RS W-Down, RS NM, FS W-Down and FS W-Up), or paired t test (G: FS NM units, L: FS NM units).

We also examined the effect of photo-inhibition of ipsilateral wM1 on the whisking-induced dynamics of wS1. The wM1 photo-inhibition led to an increase in the W-ΔAP in W-Up RS neurons (Light OFF 1.34 ± 0.14, Light ON 1.64 ± 0.21, *n* = 14 cells, *p* = 0.013, Wilcoxon signed rank test) and promoted a slight increase in the proportion of W-Up neurons by recruiting a certain fraction of (6 out of 39) NM RS neurons (Fig. 2*J-L*). While photo-inhibition of wM1 seemed to recruit some (8 out of 39) NM neurons to be W-Down neurons, the W-ΔAP in W-Down RS cells remained significantly unchanged by light (Light OFF -0.73 ± 0.05, Light ON -0.75 ± 0.06, *n* = 25 cells, *p* = 0.62, Wilcoxon signed rank test; Fig. 2*J-L*). No significant effects of wM1 photo-inhibition on FS neurons were observed (Fig. 2*J-L*). These findings suggest that wM1 activity may suppress whisking-evoked spiking in a subset of wS1 neurons. However, wM1→wS1 axons do not appear to mediate this effect, suggesting the involvement of different circuit mechanisms.

The analysis of spikes recorded with extracellular electrodes often exhibits a bias towards cells with relatively higher spike rates (Shoham et al., 2006; Barth and Poulet, 2012). This bias is especially significant in superficial layers of wS1, which contain many cells with very low spike rates that are typically overlooked by spike analyses. We therefore examined the effects of photo-inhibition of wM1→wS1 axons on subthreshold V_m_ dynamics in wS1 neurons associated with spontaneous whisking (Fig. 3). Photo-inhibition of wM1 →wS1 axons did not affect V_m_ depolarization, AP rate changes or the changes in V_m_ fluctuation upon whisking either in L2/3 or L5/6 neurons (Fig. 3*A-F*). Even in eight neurons, which exhibited an increase of AP rates by more than 0.1 Hz during whisking, photo-inhibition of wM1 → wS1 axons did not affect the magnitude of V_m_ depolarization and AP rate change upon whisking (V_m_ depolarization: Light OFF 3.9 ± 0.8 mV, Light ON 4.7 ± 0.8 mV, *p* = 0.109, Wilcoxon signed rank test; ΔAP rate: Light OFF 3.6 ± 1.8 Hz, Light ON 5.4 ± 2.8 Hz, *p* = 0.297, Wilcoxon signed rank test). Together with the results of extracellular recordings, our results thus argue against the role of wM1→wS1 signaling in the whisking-related activity of wS1, instead suggesting that the motor-sensory projection may not significantly contribute to whisking-related neuronal dynamics in wS1.

**Figure 3.**
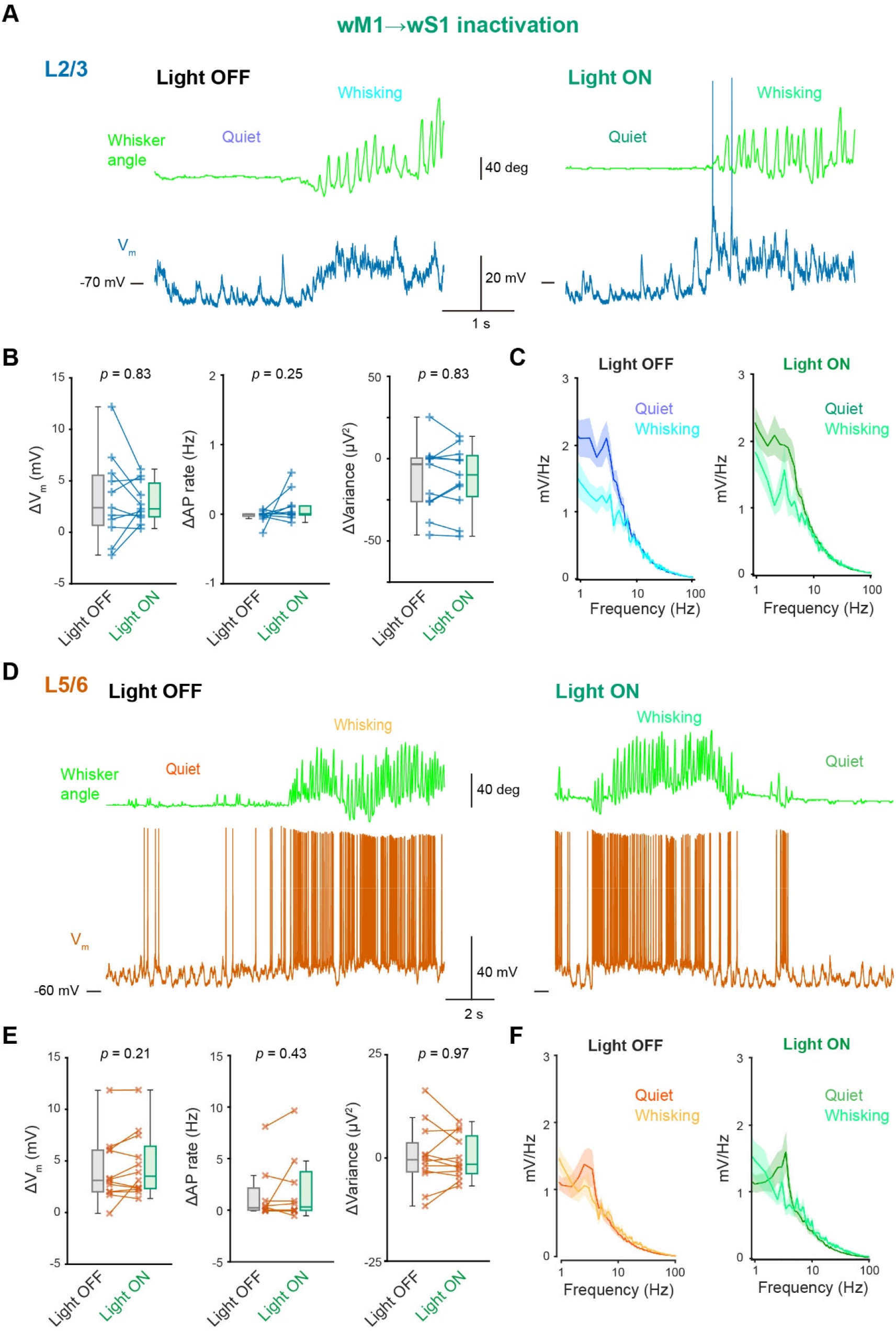
No effects of wM1→wS1 inactivation on the membrane potential of wS1 neurons. ***A***, Example membrane potential recording from a neuron at L2/3 of wS1 with (Light ON) and without (Light OFF) inactivation of the wM1→wS1 axons. ***B***, Whisking-induced changes in mean V_m_ (left), AP rate (middle), and V_m_ variance (right) of L2/3 wS1 neurons with (Light ON) and without (Light OFF) wM1→wS1 inactivation. ***C***, Averaged V_m_ FFT during quiet wakefulness and whisking (*n* = 10 cells). ***D***–***F***, Same as ***A***–***C***, but from L5/6 neurons of wS1 (*n* = 11 cells). *P*-values are indicated in the figure panels. Wilcoxon signed rank test (***B***, ***E***).

### Role of thalamocortical signaling in resting and whisking-related activity of wS1

Whisking induces a substantial increase in the AP rate of wTLM neurons (Poulet et al., 2012; Urbain et al., 2015; Petty et al., 2021). Such whisking-related activity of wTLM is known to elicit depolarization of wS1 neurons (Poulet et al., 2012) and may thus contribute to whisking-related wS1 spiking activity. However, wTLM has a broad spectrum of projections to the cerebral cortex, including wS1, wM1, and wS2 (Deschênes et al., 1998; Urbain and Deschênes, 2007b; Wimmer et al., 2010; El-Boustani et al., 2020), making unclear which projections mainly affect wS1 circuits during whisking. To clarify the role of wTLM→wS1 monosynaptic inputs in wS1 activity, we conducted photo-inhibition of the wTLM→wS1 axons using eOPN3 (Fig. 4*A,B*). During quiet wakefulness, we did not observe significant changes in the overall spontaneous AP rate of regular-spiking (RS) and fast-spiking (FS) neurons in wS1 by photo-inhibition of wTLM→wS1 axons (Fig. 4*C*). We further analyzed the effect of photo-inhibition of the wTLM→wS1 axons on the whisking-related modulation of wS1 neurons. Upon photo-inhibition, 40% (10 out of 25) of W-Up and 18.5% (12 out of 65) of W-Down RS neurons lost their responsiveness to whisking (Fig. 4*D,E*). Consistently, the averaged W-ΔAP of W-Up neurons was largely attenuated (Light OFF 1.22 ± 0.11, Light ON 0.79 ± 0.18, *n* = 25 cells, *p* = 0.0074, Wilcoxon signed rank test; Fig. 4*F*) and that of W-Down neurons was significantly increased (Light OFF -0.75 ± 0.04, Light ON -0.63 ± 0.04, *n* = 65 cells, *p* = 0.046, Wilcoxon signed rank test; Fig. 4*F*). In FS neurons, the increase in whisking-related activity of W-Up neurons was strongly inhibited by photo-inhibition (Light OFF 1.39 ± 0.18, Light ON 0.64 ± 0.25, *n* = 15 cells, *p* = 0.0083, Wilcoxon signed rank test; Fig. 4*F*). However, photo-inhibition did not affect W-Down FS neurons (Light OFF -1.29 ± 0.09, Light ON -1.12 ± 0.07, *n* = 15 cells, *p* = 0.21, Wilcoxon signed rank test; Fig. 4*F*). These results suggest that the thalamocortical axons well signal whisking to affect wS1 activity. These effects of photo-inhibition on RS and FS neurons are in good agreement with the previously proposed feed-forward inhibition circuits recruited by the thalamocortical inputs (Gabernet et al., 2005; Yu et al., 2016).

**Figure 4.**
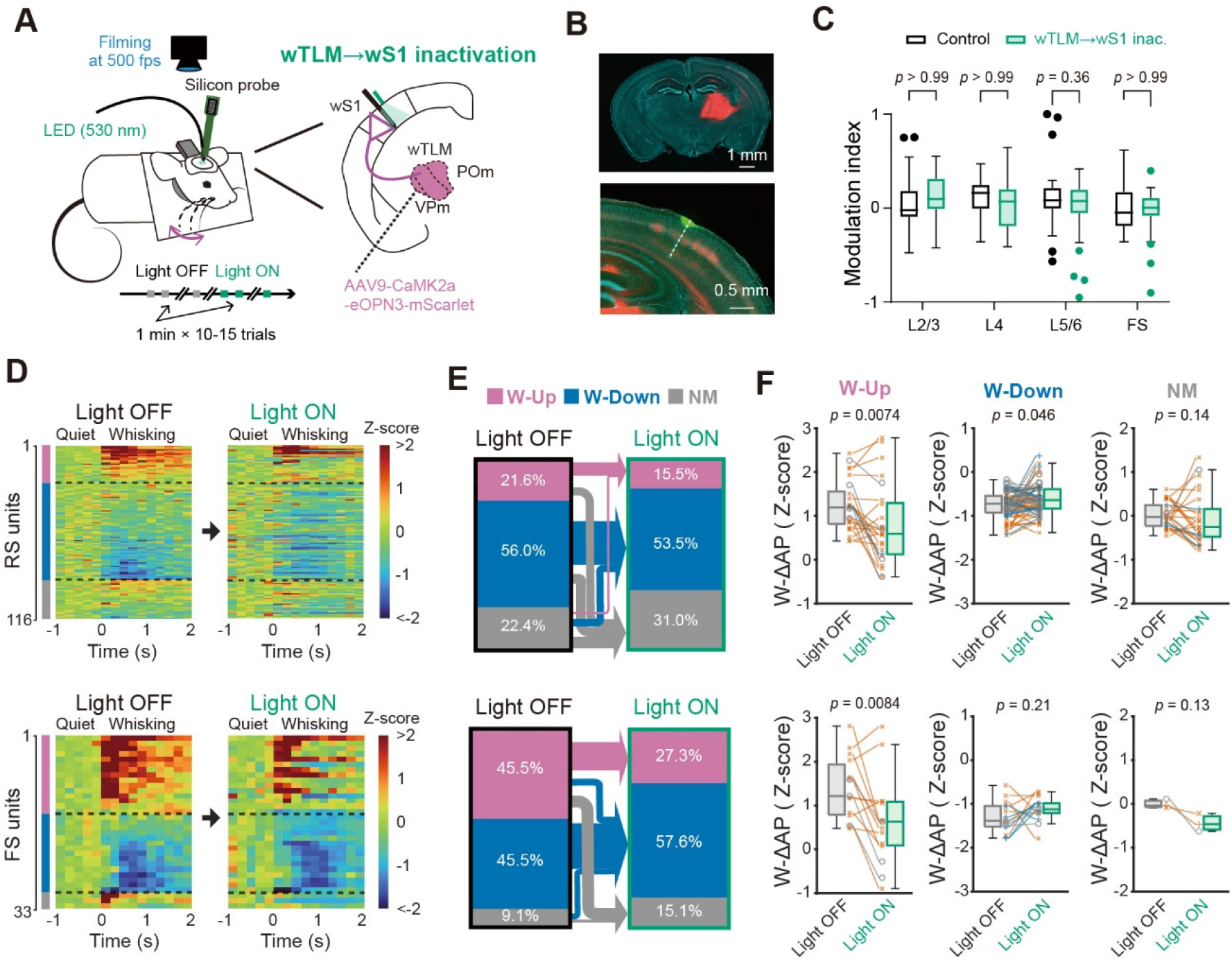
Effects of inactivation of wTLM→wS1 axons on neuronal activity in wS1. ***A***, Schematic of the experiment. ***B***, Epifluorescence image of a coronal brain section containing the wTLM injection site (top) or the wS1 recording site (bottom). The dashed line indicates the inserted electrode’s trace. Green: DiO, red: eOPN3-mScarlet, blue: DAPI. ***C***, Modulation indices of spontaneous AP rates during quiet wakefulness upon green light illumination onto wS1 in control mice (same data as in Figure 2C) or eOPN3-expressing mice (wTLM→wS1 inact., 149 cells from 3 mice). ***D***, The z-scored PSTHs of individual RS (top) and FS (bottom) units (200 ms bin size) aligned to the whisking onset with (Light ON) and without (Light OFF) inactivation of the wTLM→wS1 axons. ***E***, Proportions of whisking-modulated RS (top) or FS (bottom) units in wS1 with (Light ON) and without (Light OFF) the wTLM→wS1 inactivation. Cellular categories are the same as in Figure 2F. Arrows show transitions of the categories. ***F***, W-ΔAP (in Z-score) of RS (top) and FS (bottom) units with (Light ON) and without (Light OFF) the wTLM→wS1 inactivation. Blue lines and crosses: L2/3 units, gray lines and circles: L4 units, orange lines and saltires: L5/6 units. *P*-values are indicated in the figure panels. Bonferroni’s multiple comparison test (***C***), Wilcoxon signed rank test (***F***, RS W-Up, RS W-Down, and FS W-Up), or paired *t* test (***F***, RS NM, FS W-Down, and FS NM).

We further examined how subthreshold V_m_ dynamics of wS1 neurons can be affected by photo-inhibition of the wTLM→wS1 axons. In our whole-cell recordings, photo-inhibition of the wTLM → wS1 axons attenuated V_m_ depolarization upon whisking in both superficial and deep layers of wS1 (L2/3: Light OFF 1.3 ± 0.3 mV, Light ON -0.4 ± 0.4 mV, *n* =14 cells, *p* = 0.0031, Wilcoxon signed rank test; L5/6: Light OFF 2.7 ± 0.7 mV, Light ON 1.7 ± 0.5 mV, *n* =13 cells, *p* = 0.033, Wilcoxon signed rank test; Fig. 5*A,B,D,E*) without affecting whisking-related changes in AP rates and V_m_ variance (Fig. 5*B,E*). The photo-inhibition slightly enhanced slow-wave V_m_ oscillation during quiet wakefulness (FFT area at 1−5Hz, Light OFF 6.3 ± 0.6 mV, Light OFF 8.5 ± 0.9 mV, *n* = 27 cells, *p* = 0.030, Wilcoxon signed rank test; Fig. 5*C,F*). All these effects are similar to those previously reported after the pharmacological inhibition of wTLM (Poulet et al., 2012). Thus, a whisking-induced increase in wTLM activities directly affects wS1 neurons through direct thalamocortical connections.

**Figure 5.**
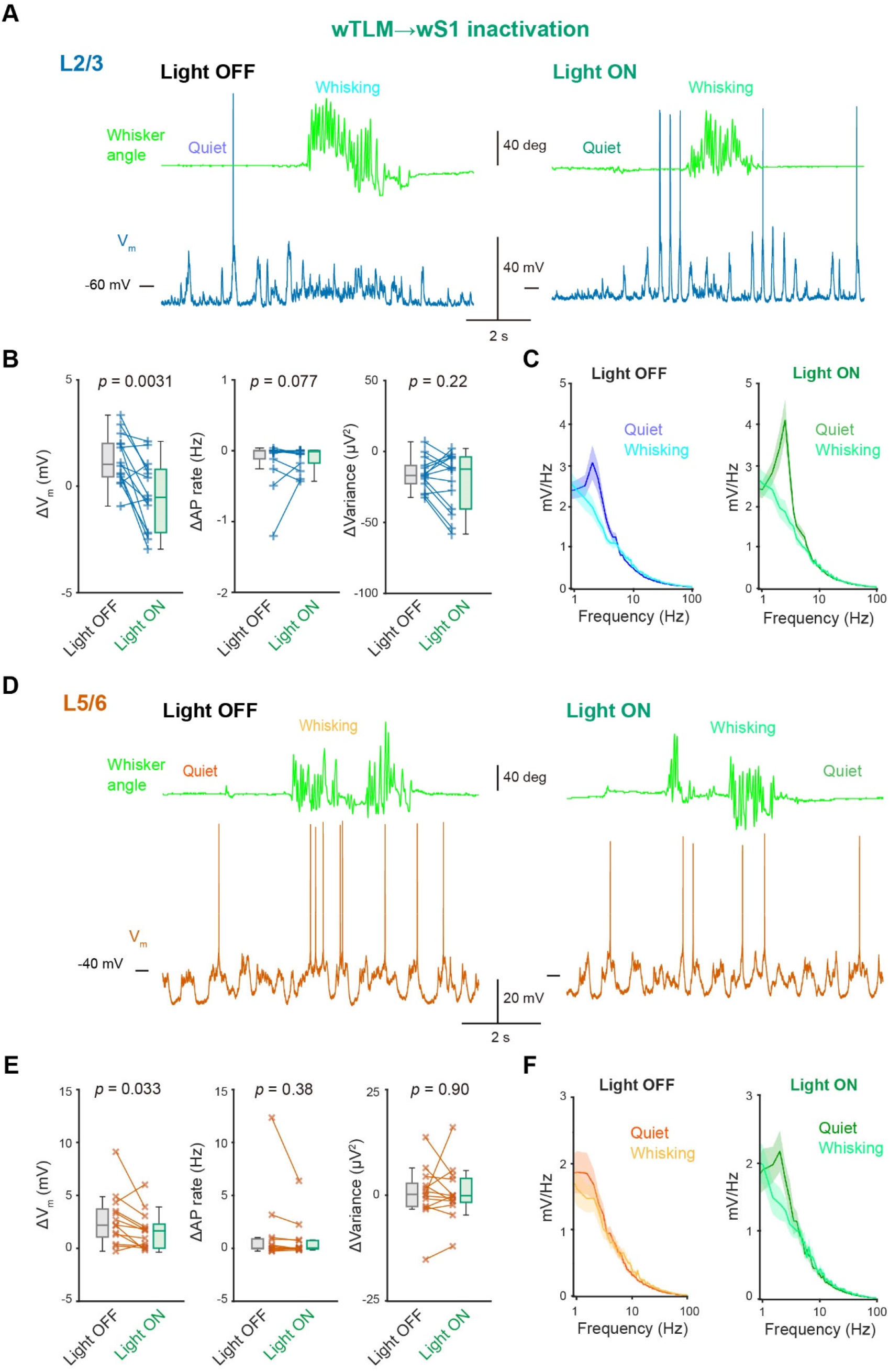
Effects of wTLM→wS1 inactivation on the membrane potential of wS1 neurons. ***A***, Example membrane potential recording from a neuron at L2/3 of wS1 with (Light ON) and without (Light OFF) inactivation of the wTLM→wS1 axons. ***B***, Whisking-induced changes in mean V_m_ (left), AP rate (middle), and V_m_ variance (right) of L2/3 wS1 neurons with (Light ON) and without (Light OFF) the wTLM→wS1 inactivation. ***C***, Averaged V_m_ FFT during quiet wakefulness and whisking (*n* = 13 cells). ***D***–***F***, Same as ***A***–***C***, but from L5/6 neurons of wS1 (*n* = 14 cells). *P*-values are indicated in the figure panels. Wilcoxon signed rank test (***B***, ***E***).

### Somatosensory feedback shapes wS1 activity in quiet and whisking states

The wS2 has a prominent motor-related activity during task performance (Chen et al., 2016; Matteucci et al., 2022), which can be signaled to wS1 through direct feedback connections (Yang et al., 2016; Minamisawa et al., 2018). We next examine the role of wS2→wS1 inputs in the whisking-related wS1 dynamics. We locally administered AAV to induce expression of eOPN3 in wS2 neurons and subsequently conducted silicon probe recordings from the C2 barrel column of ipsilateral wS1 (Fig. 6*A,B*). During quiet wakefulness, photo-inhibition of the wS2→wS1 input significantly inhibited overall spontaneous AP rates of infragranular (L5/6) RS neurons (Modulation index: mCherry [control] 0.12 ± 0.05, *n* = 36 cells, eOPN3 -0.080 ± 0.036, *n* = 65 cells, *p* = 0.0019, Bonferroni’s multiple comparisons test) but did not affect those of RS neurons in other layers and FS neurons (Fig. 6*C*). Thus, the wS2 → wS1 inputs spontaneously activate wS1 neurons in deeper layers during non-whisking, quiet states, which is consistent with more extensive innervation from wS2 to deeper layer wS1 (Minamisawa et al., 2018; Zhang and Bruno, 2019).

**Figure 6.**
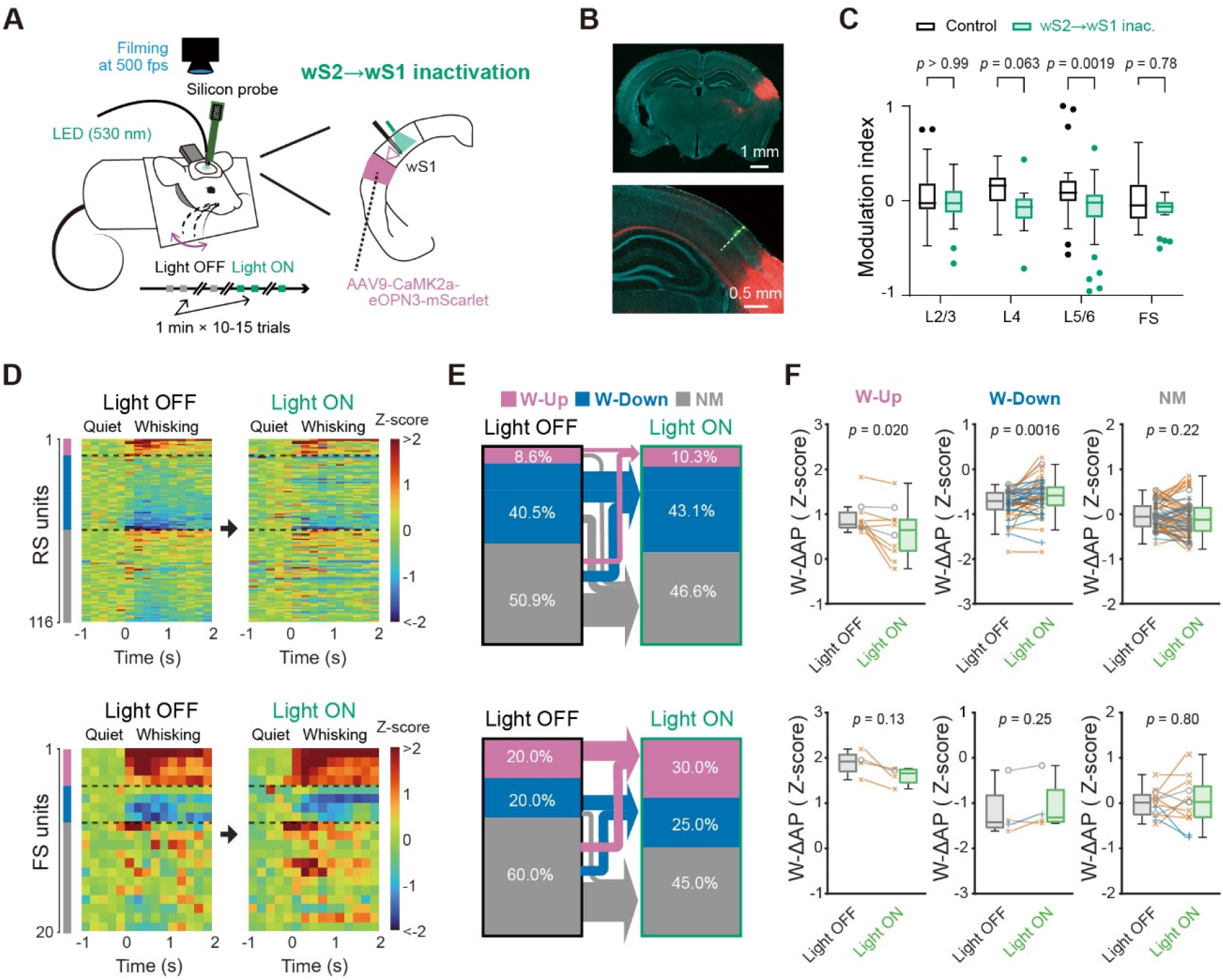
Effects of inactivation of wS2→wS1 axons on neuronal activity in wS1. ***A***, Schematic of the experiment. ***B***, Epifluorescence image of a coronal brain section containing the wS2 injection site and the wS1 recording site obtained with a lower (top) and higher (bottom) magnification. The dashed line indicates the inserted electrode’s trace. Green: DiO, red: eOPN3-mScarlet, blue: DAPI. ***C***, Modulation indices of spontaneous AP rates during quiet wakefulness upon green light illumination onto wS1 in control mice (same data as in Figure 2C) or eOPN3-expressing mice (wS2→wS1 inact., 136 cells from 3 mice). ***D***, The z-scored PSTHs of individual RS (top) and FS (bottom) units (200 ms bin size) aligned to the whisking onset with (Light ON) and without (Light OFF) inactivation of the wTLM→wS1 axons. ***E***, Proportions of whisking-modulated RS (top) or FS (bottom) units in wS1 with (Light ON) and without (Light OFF) the wTLM→wS1 inactivation. Cellular categories are the same as in Figure 2F. Arrows show transitions of the categories. ***F***, W-ΔAP (in Z-score) of RS (top) and FS (bottom) units with (Light ON) and without (Light OFF) the wTLM→wS1 inactivation. Blue lines and crosses: L2/3 units, gray lines and circles: L4 units, orange lines and saltires: L5/6 units. *P*-values are indicated in the figure panels. Bonferroni’s multiple comparison test (***C***), Wilcoxon signed rank test (***F***).

We further analyzed the effect of photo-inhibition of the wS2→wS1 axons on the whisking-related modulation of wS1 neurons. Upon photo-inhibition, 50.0% (5 out of 10) of W-Up and 21.3% (10 out of 37) of W-Down RS units lost their responsiveness to whisking (Fig. 6*D,E*). Overall, the whisking-induced changes in AP rates of W-Up and W-Down RS neurons were significantly attenuated by the wS2→wS1 photo-inhibition (W-ΔAP; W-Up: Light OFF 0.91 ± 0.11, Light ON 0.60 ± 0.17, *n* = 10 cells, *p* = 0.020, Wilcoxon signed rank test; W-Down: Light OFF -0.76 ± 0.05, Light ON -0.61 ± 0.06, *n* = 47 cells, *p* = 0.0016, Wilcoxon signed rank test; Fig. 6*F*). In contrast, the photo-inhibition did not significantly affect whisking-related modulation of FS units in wS1 (Fig. 6*D-F*).

We also performed whole-cell patch-clamp recordings from wS1 neurons to examine the effects of the wS2→wS1 photo-inhibition on V_m_ dynamics (Fig. 7). The photo-inhibition did not affect the whisking-induced changes in V_m_, AP rate, V_m_ variance, and V_m_ fluctuation (Fig. 7*B,C*) in neurons at L2/3. At L5/6, the wS2→wS1 photo-inhibition apparently reduced the whisking-induced AP rate changes (Light OFF 1.0 ± 0.7 Hz, Light ON 0.1 ± 0.1 Hz, *n* = 12 cells, *p* = 0.037; Fig. 7*E*), but its effect on the whisking-induced V_m_ depolarization, V_m_ variance, and slow-wave V_m_ fluctuation was not evident (Fig. 7*E,F*). In eight neurons that exhibited an increase of AP rates by more than 0.1 Hz during whisking, significant suppression of whisking-related depolarization and spiking was observed with the wS2 → wS1 photo-inhibition (V_m_ depolarization: Light OFF, 4.2 ± 1.1 mV, Light ON, 2.6 ± 0.9 mV, *p* = 0.0156, Wilcoxon signed rank test; ΔAP rate: Light OFF, 2.0 ± 0.9 Hz, Light ON, 0.4 ± 0.1 Hz, *p* = 0.0078, Wilcoxon signed rank test; Fig. 8). Our results thus indicate that the somatosensory feedback connections from wS2 to wS1 enhance the whisking responsiveness of a subset of wS1 neurons and create the whisking-related dynamics in wS1.

**Figure 7.**
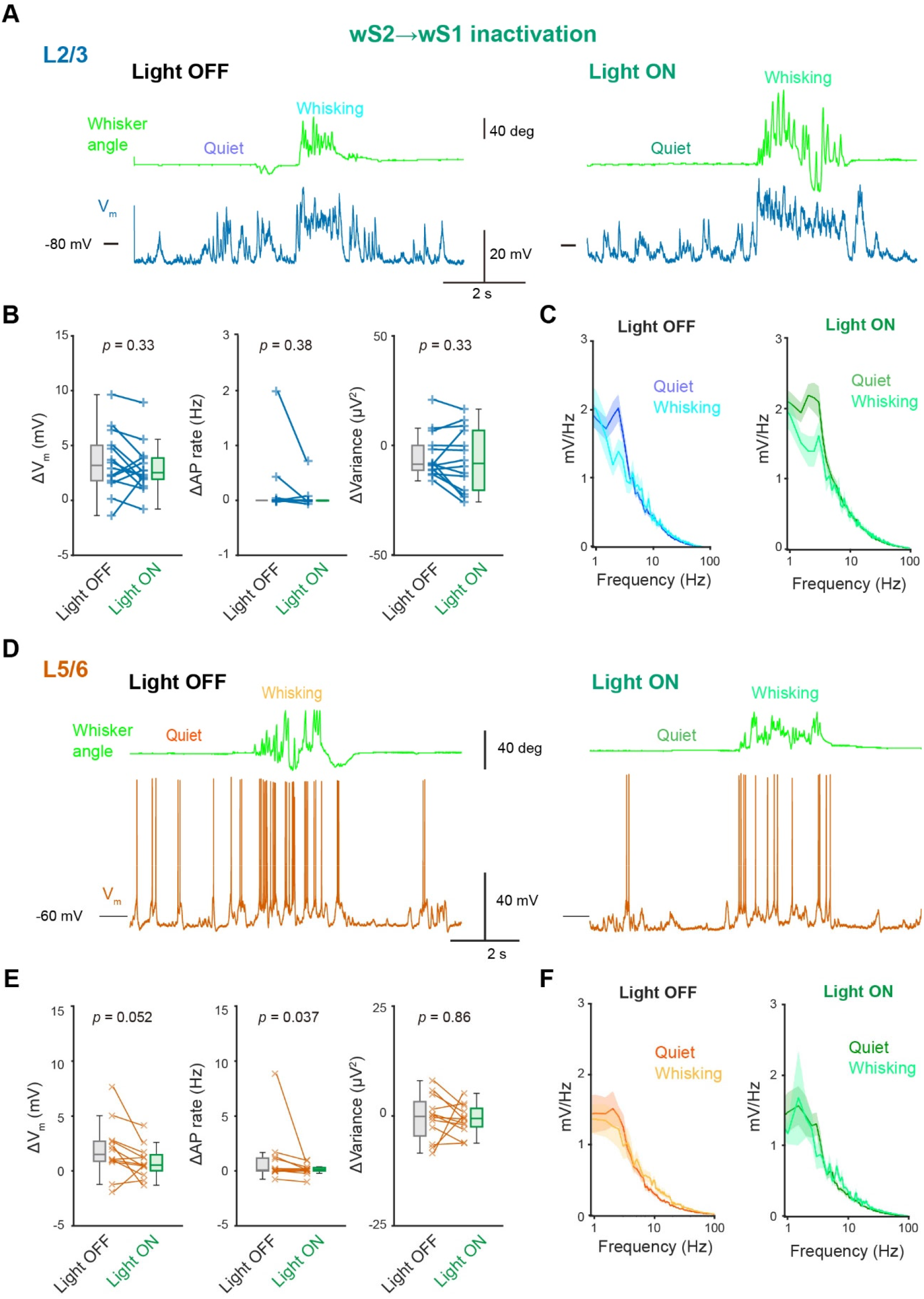
Effects of wS2→wS1 inactivation on the membrane potential of wS1 neurons. ***A***, Example membrane potential recording from a neuron at L2/3 of wS1 with (Light ON) and without (Light OFF) inactivation of the wS2→wS1 axons. ***B***, Whisking-induced changes in mean V_m_ (left), AP rate (middle), and V_m_ variance (right) of L2/3 wS1 neurons with (Light ON) and without (Light OFF) the wS2→wS1 inactivation. ***C***, Averaged V_m_ FFT during quiet wakefulness and whisking (*n* = 12 cells). ***D***–***F***, Same as ***A***–***C***, but from L5/6 neurons of wS1 (*n* = 10 cells). *P*-values are indicated in the figure panels. Wilcoxon signed rank test (***B***, ***E***).

**Figure 8.**
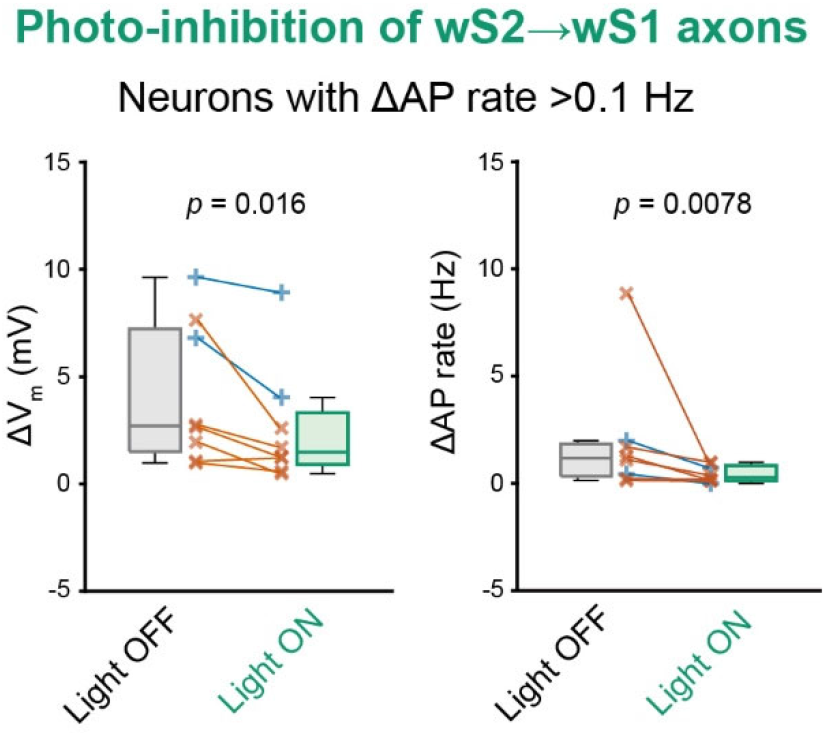
Effects of wS2→wS1 inactivation on the membrane potential of the wS1 neurons positively modulated by whisking. Whisking-induced changes in mean V_m_ (left), and AP rate (right) and V_m_ variance (right) of wS1 neurons with (Light ON) and without (Light OFF) the wS2→wS1 inactivation. Lines indicate individual data from L2/3 (blue) or L5/6 (orange). *P*-values are indicated in the figure panels. Wilcoxon signed rank test.

## Discussion

Using eOPN3-mediated photo-inhibition of synaptic transmission, we investigated the role of specific inputs from wM1, wTLM, and wS2 to wS1 in the whisking-related modulation of supra- and sub-threshold V_m_ of wS1 neurons. We found that wTLM→wS1 and wS2→wS1 projections to wS1 significantly shape the whisking-related modulation of spike rates of wS1 neurons. These two pathways also enhance the whisking-induced depolarization of wS1 neurons. In contrast, wM1 appears to have little overall influence on the whisking-related modulation of wS1 neurons, despite its rich innervation and contribution to spontaneous AP rates in wS1 deeper layer neurons.

Neurons in wM1 represent various aspects of whisker movements (Hill et al., 2011; Friedman et al., 2012; Gerdjikov et al., 2013; Sreenivasan et al., 2016), and stimulation of wM1 neurons initiates rhythmic whisking (Ferezou et al., 2007; Matyas et al., 2010; Sreenivasan et al., 2015; Sreenivasan et al., 2016). The wM1 neurons have extensive axonal projections to L1 and L5 at wS1, with monosynaptic connections to wS1 neurons (Petreanu et al., 2009; Kinnischtzke et al., 2014). Optogenetic stimulation of wM1 axons in wS1 *in vivo* causes state changes in wS1 neurons, similar to whisking-induced ones (Zagha et al., 2013). Among wS1 neurons, GABAergic neurons expressing vasoactive intestinal peptide-expressing (VIP) receive the most robust excitatory inputs from wM1 (Lee et al., 2013; Naskar et al., 2021). Excitation of VIP neurons is known to disinhibit nearby excitatory pyramidal neurons, mostly at distal dendrites, by inhibiting somatostatin-expressing (SST) GABAergic neurons (Lee et al., 2013; Pfeffer et al., 2013; Pi et al., 2013; Zhang et al., 2014). Moreover, regional inhibition of wM1 causes insensitivity of VIP and SST neurons in wS1 to whisking (Lee et al., 2013) and largely attenuates dendritic touch responses of wS1 L5 neurons (Xu et al., 2012). Therefore, it has long been hypothesized that wM1 neurons would affect wS1 neurons through exciting VIP neurons during whisking, recruiting the disinhibitory circuit. Nevertheless, our results provide little evidence of the role of wM1→wS1 inputs in whisking-induced modulation in the AP rates and V_m_ dynamics of wS1 neurons. Instead, our results suggest that wM1→wS1 inputs increase spontaneous AP rates of wS1 neurons during quiet wakefulness.

In the primary auditory cortex (A1), like in wS1, optogenetic stimulation of the top-down projections from the frontal motor cortex to A1 induces depolarization and the reduction of slow-wave V_m_ oscillation in A1 neurons as seen upon body movements (Schneider et al., 2014). The disinhibitory VIP-SST circuit exists in A1 as well (Pi et al., 2013). However, the disinhibitory mechanism seems not to operate for the motor-related modulation of A1 neurons (Yavorska and Wehr, 2021). Primary visual cortex (V1) also implicates the disinhibitory circuit involving VIP and SST neurons, which is recruited mainly by cholinergic inputs, controlling the gain of sensory responses (Fu et al., 2014). In wS1, cholinergic axons from the basal forebrain are activated during whisking and release acetylcholine to excite VIP neurons even after shutting down local glutamatergic synaptic transmission (Eggermann et al., 2014; Gasselin et al., 2021). Taken together, the top-down wM1→wS1 inputs might not be the primary mechanism for generating whisking-induced state changes in wS1 neurons. Rather, cholinergic inputs may have a major role. Future investigations might require exploring the interactions between neuromodulatory systems and whisker sensory-motor inputs and their potential role in state-dependent modulation and sensory processing.

During episodes of whisking, neurons within the wTLM receive excitatory synaptic inputs from sources yet to be identified, which enhance their firing rates (Moore et al., 2015; Urbain et al., 2015). Our data illustrates that inactivation of the wTLM→wS1 projections amplifies slow-wave V_m_ oscillations and decreases the depolarization of wS1 neurons during whisking. These observations echo those made in a previous study that employed regional wTLM inhibition (Poulet et al., 2012), supporting a hypothesis that wTLM’s role in the whisking-related state changes within wS1 is, at least partially, mediated through direct synaptic inputs to wS1. In addition, our results highlight the potential influence of wS2→wS1 feedback signaling in modulating wS1 activity during whisking. The effect of the photo-inhibition of wS2→wS1 inputs on spontaneous AP rates and whisking-related depolarization was primarily observable in the deeper layers (L5/6) of wS1. This result aligns with anatomical evidence showing more extensive innervation of wS2 axons within these deeper layers than L2/3 (Minamisawa et al., 2018; Zhang and Bruno, 2019). Given that wS2 receives excitatory synaptic inputs from wTLM, it is plausible that wS2 serves as a relay for whisking-related wTLM activity towards wS1 (El-Boustani et al., 2020). Furthermore, wS2 also receives top-down inputs from wM2, which may play a critical role in transmitting motor-related signals to wS1 via wS2 (Matteucci et al., 2022).

Our whole-cell recordings revealed at least part of the mechanisms underlying whisking-related depolarization of wS1 cells, accounting for the presence of W-Up cells in wS1. However, while approximately half of the extracellularly recorded units were identified as W-Down cells, the intracellular mechanisms underpinning their existence remain elusive. Despite the activities of these W-Down units seemingly being shaped by wTLM→wS1 and wS2→wS1 projections, our whole-cell recordings failed to capture these mechanisms - potentially due to lower spontaneous AP rates and shorter recording duration. This reinforces the need for technological advancements to secure sufficient spontaneous whisking epochs for a reliable comparison of AP rates.

Although our analysis primarily focuses on wS1 activity during spontaneous behaviors, the role of specific signaling pathways to wS1 can be modulated in mice engaged in cognitive or sensorimotor tasks. For instance, following the learning of a simple association between whisker stimulation and water reward availability in a whisker-detection task, a subset of wS1 neurons, particularly those projecting to wS2, show sensitivity to goal-directed licking behavior and undergo depolarization upon licking (Yamashita and Petersen, 2016). In line with this, the neurons within wS2 have been observed to encode goal-directed licking more accurately than wS1 and another task-related brain region, wM2, suggesting a potential enhancement of wS2→wS1 signaling during goal-directed movement (Matteucci et al., 2022). Furthermore, the wM1 neurons projecting to wS1 also encode a variety of task- and motor-related information (Petreanu et al., 2012). Future studies involving task-performing mice will thus be crucial in further elucidating the role of specific long-range inputs to wS1 in shaping neuronal activity in wS1 during task-related movements.

## Author contributions

M.K. and T.Y. designed research; M.K. performed research; M.K., K.H., M.O., and W.L. analyzed data; M.K. wrote the first draft of the paper; M.K. and T.Y. wrote the paper.

## Acknowledgements

We thank Dr. Carl Petersen for discussion; E. Imoto and M. Jin for technical assistance. This work was supported by JST FOREST Program (JPMJFR204H), JSPS KAKENHI grants (21H00215, 21K19315, 23H04366, and 23H04685) to T.Y.; The Naito Foundation to T.Y.; Takeda Science Foundation to T.Y.; Research Foundation for the Electrotechnology of Chubu to T.Y. M.K. was supported by the Nagoya University CIBoG WISE program of MEXT.

## References

1. Barth AL, Poulet JF (2012) Experimental evidence for sparse firing in the neocortex. Trends Neurosci 35:345–355.

2. Benjamini Y, Hochberg Y (1995) Controlling the False Discovery Rate: A Practical and Powerful Approach to Multiple Testing. Journal of the Royal Statistical Society: Series B (Methodological) 57:289–300.

3. Chen JL, Voigt FF, Javadzadeh M, Krueppel R, Helmchen F (2016) Long-range population dynamics of anatomically defined neocortical networks. Elife 5.

4. Cheung JA, Maire P, Kim J, Lee K, Flynn G, Hires SA (2020) Independent representations of self-motion and object location in barrel cortex output. PLoS Biol 18:e3000882.

5. Crochet S, Petersen CCH (2006) Correlating whisker behavior with membrane potential in barrel cortex of awake mice. Nature Neuroscience 9:608–610.

6. de Kock CP, Sakmann B (2009) Spiking in primary somatosensory cortex during natural whisking in awake head-restrained rats is cell-type specific. Proc Natl Acad Sci U S A 106:16446–16450.

7. Deschênes M, Veinante P, Zhang ZW (1998) The organization of corticothalamic projections: reciprocity versus parity. Brain Res Brain Res Rev 28:286–308.

8. Eggermann E, Kremer Y, Crochet S, Petersen CCH (2014) Cholinergic signals in mouse barrel cortex during active whisker sensing. Cell Rep 9:1654–1660.

9. El-Boustani S, Sermet BS, Foustoukos G, Oram TB, Yizhar O, Petersen CCH (2020) Anatomically and functionally distinct thalamocortical inputs to primary and secondary mouse whisker somatosensory cortices. Nat Commun 11:3342.

10. Fee MS, Mitra PP, Kleinfeld D (1997) Central versus peripheral determinants of patterned spike activity in rat vibrissa cortex during whisking. J Neurophysiol 78:1144–1149.

11. Ferezou I, Bolea S, Petersen CC (2006) Visualizing the cortical representation of whisker touch: voltage-sensitive dye imaging in freely moving mice. Neuron 50:617–629.

12. Ferezou I, Haiss F, Gentet LJ, Aronoff R, Weber B, Petersen CC (2007) Spatiotemporal dynamics of cortical sensorimotor integration in behaving mice. Neuron 56:907–923.

13. Freeman JA, Nicholson C (1975) Experimental optimization of current source-density technique for anuran cerebellum. J Neurophysiol 38:369–382.

14. Friedman WA, Zeigler HP, Keller A (2012) Vibrissae motor cortex unit activity during whisking. J Neurophysiol 107:551–563.

15. Fu Y, Tucciarone JM, Espinosa JS, Sheng N, Darcy DP, Nicoll RA, Huang ZJ, Stryker MP (2014) A cortical circuit for gain control by behavioral state. Cell 156:1139–1152.

16. Gabernet L, Jadhav SP, Feldman DE, Carandini M, Scanziani M (2005) Somatosensory integration controlled by dynamic thalamocortical feed-forward inhibition. Neuron 48:315–327.

17. Gallant JL, Connor CE, Van Essen DC (1998) Neural activity in areas V1, V2 and V4 during free viewing of natural scenes compared to controlled viewing. Neuroreport 9:2153–2158.

18. Gasselin C, Hohl B, Vernet A, Crochet S, Petersen CCH (2021) Cell-type-specific nicotinic input disinhibits mouse barrel cortex during active sensing. Neuron 109:778–787.e773.

19. Gerdjikov TV, Haiss F, Rodriguez-Sierra OE, Schwarz C (2013) Rhythmic whisking area (RW) in rat primary motor cortex: an internal monitor of movement-related signals? J Neurosci 33:14193–14204.

20. Hill DN, Curtis JC, Moore JD, Kleinfeld D (2011) Primary motor cortex reports efferent control of vibrissa motion on multiple timescales. Neuron 72:344–356.

21. Hubel DH, Wiesel TN (1962) Receptive fields, binocular interaction and functional architecture in the cat’s visual cortex. J Physiol 160:106–154.

22. Katsuki Y, Watanabe T, Maruyama N (1959) Activity of auditory neurons in upper levels of brain of cat. J Neurophysiol 22:343–359.

23. Kinnischtzke AK, Simons DJ, Fanselow EE (2014) Motor cortex broadly engages excitatory and inhibitory neurons in somatosensory barrel cortex. Cereb Cortex 24:2237–2248.

24. Lee S, Kruglikov I, Huang ZJ, Fishell G, Rudy B (2013) A disinhibitory circuit mediates motor integration in the somatosensory cortex. Nature Neuroscience 16:1662–1670.

25. Lefort S, Tomm C, Floyd Sarria JC, Petersen CC (2009) The excitatory neuronal network of the C2 barrel column in mouse primary somatosensory cortex. Neuron 61:301–316.

26. Mahn M et al. (2021) Efficient optogenetic silencing of neurotransmitter release with a mosquito rhodopsin. Neuron 109:1621–1635.e1628.

27. Matteucci G, Guyoton M, Mayrhofer JM, Auffret M, Foustoukos G, Petersen CCH, El-Boustani S (2022) Cortical sensory processing across motivational states during goal-directed behavior. Neuron 110:4176–4193.e4110.

28. Matyas F, Sreenivasan V, Marbach F, Wacongne C, Barsy B, Mateo C, Aronoff R, Petersen CCH (2010) Motor Control by Sensory Cortex. Science 330:1240–1243.

29. Miguel P-V, Mikhail AL, Michael CW, Miguel ALN (2013) Simultaneous Top-down Modulation of the Primary Somatosensory Cortex and Thalamic Nuclei during Active Tactile Discrimination. The Journal of Neuroscience 33:4076.

30. Minamisawa G, Kwon SE, Chevée M, Brown SP, O’Connor DH (2018) A Non-canonical Feedback Circuit for Rapid Interactions between Somatosensory Cortices. Cell Rep 23:2718–2731.e2716.

31. Moore JD, Mercer Lindsay N, Deschênes M, Kleinfeld D (2015) Vibrissa Self-Motion and Touch Are Reliably Encoded along the Same Somatosensory Pathway from Brainstem through Thalamus. PLoS Biol 13:e1002253.

32. Mountcastle VB, Davies PW, Berman AL (1957) Response properties of neurons of cat’s somatic sensory cortex to peripheral stimuli. J Neurophysiol 20:374–407.

33. Naskar S, Qi J, Pereira F, Gerfen CR, Lee S (2021) Cell-type-specific recruitment of GABAergic interneurons in the primary somatosensory cortex by long-range inputs. Cell Reports 34.

34. Niell CM, Stryker MP (2010) Modulation of visual responses by behavioral state in mouse visual cortex. Neuron 65:472–479.

35. Otchy TM, Wolff SBE, Rhee JY, Pehlevan C, Kawai R, Kempf A, Gobes SMH, Ölveczky BP (2015) Acute off-target effects of neural circuit manipulations. Nature 528:358–363.

36. Pala A, Stanley GB (2022) Ipsilateral Stimulus Encoding in Primary and Secondary Somatosensory Cortex of Awake Mice. J Neurosci 42:2701–2715.

37. Petreanu L, Mao T, Sternson SM, Svoboda K (2009) The subcellular organization of neocortical excitatory connections. Nature 457:1142–1145.

38. Petreanu L, Gutnisky DA, Huber D, Xu NL, O’Connor DH, Tian L, Looger L, Svoboda K (2012) Activity in motor-sensory projections reveals distributed coding in somatosensation. Nature 489:299–303.

39. Petty GH, Kinnischtzke AK, Hong YK, Bruno RM (2021) Effects of arousal and movement on secondary somatosensory and visual thalamus. eLife 10:e67611.

40. Pfeffer CK, Xue M, He M, Huang ZJ, Scanziani M (2013) Inhibition of inhibition in visual cortex: the logic of connections between molecularly distinct interneurons. Nat Neurosci 16:1068–1076.

41. Pi H-J, Hangya B, Kvitsiani D, Sanders JI, Huang ZJ, Kepecs A (2013) Cortical interneurons that specialize in disinhibitory control. Nature 503:521–524.

42. Polack PO, Friedman J, Golshani P (2013) Cellular mechanisms of brain state-dependent gain modulation in visual cortex. Nat Neurosci 16:1331–1339.

43. Poulet JF, Petersen CC (2008) Internal brain state regulates membrane potential synchrony in barrel cortex of behaving mice. Nature 454:881–885.

44. Poulet JFA, Fernandez LMJ, Crochet S, Petersen CCH (2012) Thalamic control of cortical states. Nature Neuroscience 15:370–372.

45. Schneider DM, Nelson A, Mooney R (2014) A synaptic and circuit basis for corollary discharge in the auditory cortex. Nature 513:189–194.

46. Shoham S, O’Connor DH, Segev R (2006) How silent is the brain: is there a “dark matter” problem in neuroscience? J Comp Physiol A Neuroethol Sens Neural Behav Physiol 192:777–784.

47. Sreenivasan V, Karmakar K, Rijli FM, Petersen CC (2015) Parallel pathways from motor and somatosensory cortex for controlling whisker movements in mice. Eur J Neurosci 41:354–367.

48. Sreenivasan V, Esmaeili V, Kiritani T, Galan K, Crochet S, Petersen CCH (2016) Movement Initiation Signals in Mouse Whisker Motor Cortex. Neuron 92:1368–1382.

49. Suryadeep D, Dawn MA, Shane RC (2022) State-Dependent Modulation of Activity in Distinct Layer 6 Corticothalamic Neurons in Barrel Cortex of Awake Mice. The Journal of Neuroscience 42:6551.

50. Urbain N, Deschênes M (2007a) Motor Cortex Gates Vibrissal Responses in a Thalamocortical Projection Pathway. Neuron 56:714–725.

51. Urbain N, Deschênes M (2007b) A New Thalamic Pathway of Vibrissal Information Modulated by the Motor Cortex. The Journal of Neuroscience 27:12407.

52. Urbain N, Salin PA, Libourel PA, Comte JC, Gentet LJ, Petersen CCH (2015) Whisking-Related Changes in Neuronal Firing and Membrane Potential Dynamics in the Somatosensory Thalamus of Awake Mice. Cell Rep 13:647–656.

53. Wang HY, Eguchi K, Yamashita T, Takahashi T (2020) Frequency-Dependent Block of Excitatory Neurotransmission by Isoflurane via Dual Presynaptic Mechanisms. J Neurosci 40:4103–4115.

54. Wimmer VC, Bruno RM, de Kock CP, Kuner T, Sakmann B (2010) Dimensions of a projection column and architecture of VPM and POm axons in rat vibrissal cortex. Cereb Cortex 20:2265–2276.

55. Xu NL, Harnett MT, Williams SR, Huber D, O’Connor DH, Svoboda K, Magee JC (2012) Nonlinear dendritic integration of sensory and motor input during an active sensing task. Nature 492:247–251.

56. Yamashita T, Petersen CCH (2016) Target-specific membrane potential dynamics of neocortical projection neurons during goal-directed behavior. eLife 5:e15798.

57. Yamashita T, Pala A, Pedrido L, Kremer Y, Welker E, Petersen Carl CH (2013) Membrane Potential Dynamics of Neocortical Projection Neurons Driving Target-Specific Signals. Neuron 80:1477–1490.

58. Yang H, Kwon SE, Severson KS, O’Connor DH (2016) Origins of choice-related activity in mouse somatosensory cortex. Nat Neurosci 19:127–134.

59. Yavorska I, Wehr M (2021) Effects of Locomotion in Auditory Cortex Are Not Mediated by the VIP Network. Front Neural Circuits 15:618881.

60. Yu J, Gutnisky DA, Hires SA, Svoboda K (2016) Layer 4 fast-spiking interneurons filter thalamocortical signals during active somatosensation. Nat Neurosci 19:1647–1657.

61. Zagha E, Casale Amanda E, Sachdev Robert NS, McGinley Matthew J, McCormick David A (2013) Motor Cortex Feedback Influences Sensory Processing by Modulating Network State. Neuron 79:567–578.

62. Zhang S, Xu M, Kamigaki T, Hoang Do JP, Chang W-C, Jenvay S, Miyamichi K, Luo L, Dan Y (2014) Long-range and local circuits for top-down modulation of visual cortex processing. Science 345:660–665.

63. Zhang W, Bruno RM (2019) High-order thalamic inputs to primary somatosensory cortex are stronger and longer lasting than cortical inputs. eLife 8:e44158.

64. Zhang Z, Zagha E (2023) Motor cortex gates distractor stimulus encoding in sensory cortex. Nature Communications 14:2097.

65. Zhou M, Liang F, Xiong XR, Li L, Li H, Xiao Z, Tao HW, Zhang LI (2014) Scaling down of balanced excitation and inhibition by active behavioral states in auditory cortex. Nat Neurosci 17:841–850.

